# Anthropogenic remediation of heavy metals selects against natural microbial remediation

**DOI:** 10.1101/441873

**Authors:** Elze Hesse, Daniel Padfield, Florian Bayer, Eleanor M. van Veen, Christopher G. Bryan, Angus Buckling

## Abstract

In an era of unprecedented environmental change, there have been increasing ecological and global public health concerns associated with exposure to anthropogenic pollutants. While there is a pressing need to remediate polluted ecosystems, human intervention strategies might unwittingly oppose selection for natural detoxification, which is primarily carried out by microbes. We test this possibility in the context of a ubiquitous chemical remediation strategy aimed at targeting toxic metal pollution: the addition of lime-containing materials. Here we show that raising pH by liming decreased the availability of toxic metals in acidic mine-degraded soils, but as a consequence selected against microbial taxa that naturally remediate soil through the production of metal-scavenging siderophores. Understanding the ecological and evolutionary consequences of human intervention on key traits is crucial for the engineering of evolutionary resilient microbial communities, having important implications for human health and biotechnology.

## Introduction

Intervention strategies aimed at improving human health, agriculture, biotechnology and the environment inevitably impact microbial communities; even in situations where microorganisms are not the direct targets of intervention. Given the potential of microbial populations and communities to rapidly respond to any environmental change (Bissett et al. 2013; Green et al. 2008; Lau and Lennon 2012; Li et al. 2014; Zogg et al. 1997), it is crucial to consider both the short- and long-term effects resulting from ecological species sorting and adaptation. Of particular concern is if longer-term responses reduce the efficacy of intervention or result in negative consequences. This might occur if the intervention opposes the naturally human-beneficial characteristics of microbes. This study investigates such a scenario in the context of raising pH using lime-containing materials, a common intervention practice aimed at reducing heavy metal toxicity (Wuana and Okieimen 2011).

Heavy metals (metals and metalloids with a density above 5 g^-1^ cm^3^) are ubiquitous components of the Earth’s crust (Nriagu and Pacyna 1988). As a result of soaring demands for minerals (Douce 2016), large parts of the world are currently mined for valuable mineral deposits, leaving a legacy of untreated mining waste (Dudka and Adriano 1997). In addition, agricultural practices such as application of sewage sludge and phosphate fertilisers have led to increased heavy metal concentrations in the environment (Ma et al. 2018; Puschenreiter et al. 2005). Heavy metals typically persist for a long time after their introduction (Adriano 2001) and can adversely affect human, plant and wild-life at high concentrations (Tchounwou et al. 2012). Consequently, there is increasing interest in remediating metal-contaminated environments. Acidic conditions often prevail in mine-degraded sites, and the bioavailability of many heavy metals is increased under such conditions (Wuana and Okieimen 2011). Hence, lime-containing materials are commonly applied to neutralize soil pH (Edmeades and Ridley 2003), immobilize heavy metals (Sheoran et al. 2010) and thereby facilitate natural regeneration (Hamilton et al. 2007; Paradelo et al. 2015; Wuana and Okieimen 2011).

Microbial communities inhabiting contaminated soils have evolved various resistance mechanisms, including sequestration, efflux and extracellular chelation (Bruins et al. 2000; Nies 1999; Valls and De Lorenzo 2002). Crucially, some of these mechanisms – notably chelation – also act to remediate the environment. Much of this chelation is carried out by siderophores – low molecular weight high-affinity iron chelators that are synthesized and secreted by many microorganisms in response to iron deprivation (Hider and Kong 2010). While the canonical function of siderophores is iron scavenging, these secreted molecules also bind to other metals, thereby preventing their uptake into cells and rendering the environment less toxic (Braud et al. 2009; Schalk et al. 2011). Our recent experimental work has shown that siderophores provide across-species protective benefits against toxic metals (O’Brien et al. 2018) and that siderophore-producing microbial taxa are selectively favoured in metal-contaminated soils (Hesse, O’Brien, et al. 2018). It therefore follows that raising pH will select against this natural decontamination process carried out by the microbes.

We used an experimental evolution approach to determine how liming of acidic mine-degraded soils influences microbial community function (i.e. through the production of metal-chelating siderophores) and composition. We collected thirty distinct soil samples in a historical mining area to determine whether liming has a consistent effect across soils varying widely in their initial pH, metal concentrations and community compositions (Hesse, O’Brien, et al. 2018). Using a paired design, we subjected each soil to two different selection regimes by incubating soil microcosms for 12 weeks with and without the addition of hydrated lime. Soil characteristics, siderophore production and community composition were quantified before and after experimental manipulation. Our results demonstrate that liming opposes natural selection by favouring microbial taxa that produce little or no siderophores, leading to a net decrease in community-wide siderophore production.

## Results

### Liming reduces soil acidity and non-iron metal availability

Soils across the mining-contaminated gradient varied widely in their pH (lmer: estimated standard deviation of community origin random effect = 0.97), being predominantly acidic before experimental manipulation (Fig. 1a). Liming had the desired effect of raising pH (treatment effect*: χ^2^_2_* = 64.85, *P* < 0.001; Fig. 1b), such that pH was significantly greater in lime-treated soils 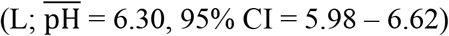 compared to ancestral 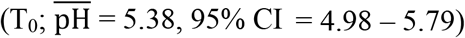 and control 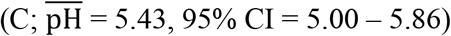 soils (Tukey contrasts for C – T_0_: *z* = 0.53 and *P =* 0.86, L – T_0_: *z* = 9.46 and *P* < 0.001 and L – C: *z* = 8.93 and *P* < 0.001).

**Figure 1.**
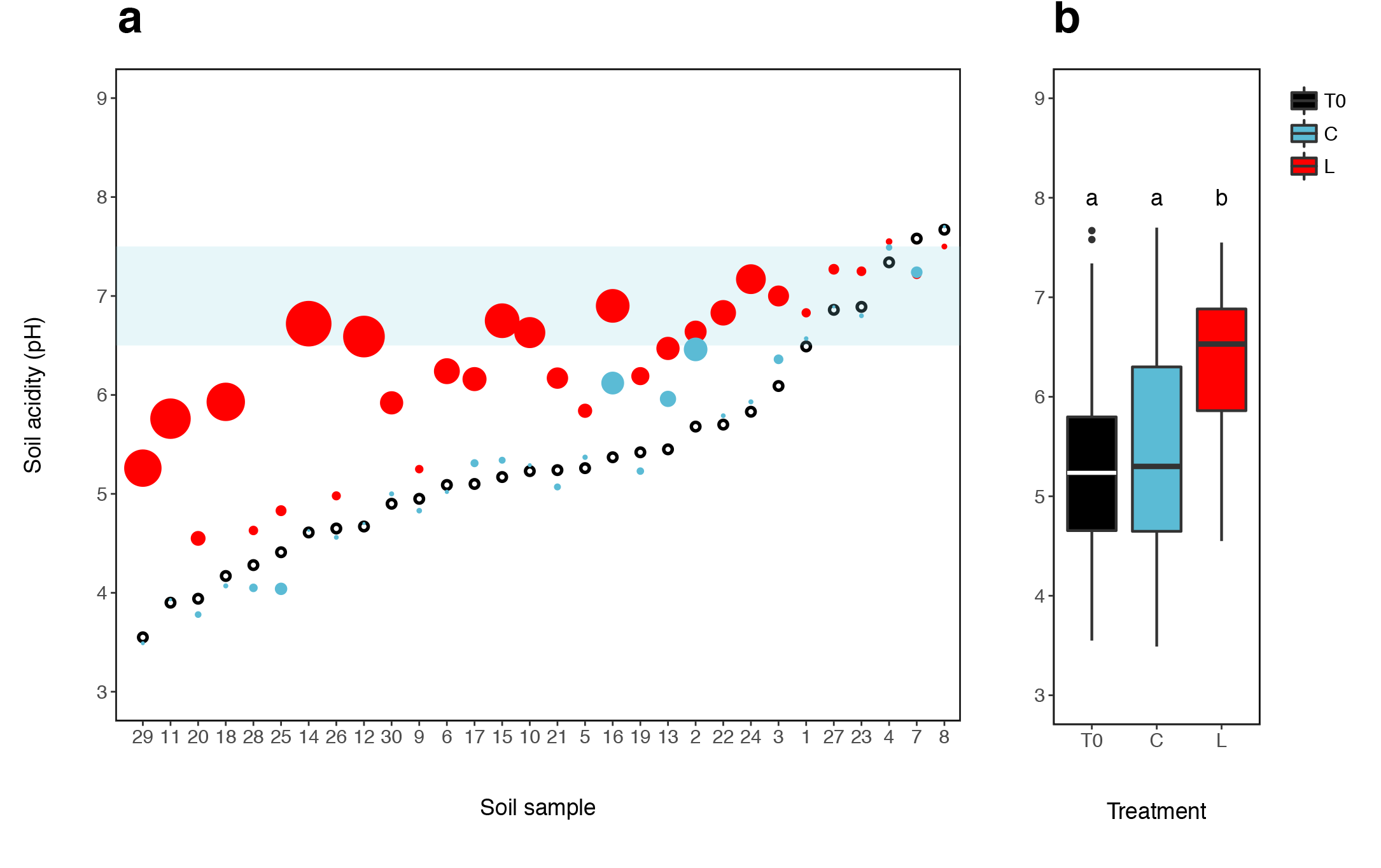
Variation in soil acidity (pH) in a historical mining site. Panel **a** shows that soils were predominantly acidic (T0, black symbols), falling well below the blue shaded area of pH neutrality. Symbol size indicates the extent to which the lime (L, red symbols) and control (C, blue symbols) treatments altered initial soil acidity (T_0_). Note that the effect of lime addition is greater for initially more acidic soils. **(b)** Box whisker plot showing a reduction in soil acidity in response to lime addition. Boxes depict the upper and lower quartiles of the treatment-specific raw data with the centre line showing the median and the whiskers providing a measure of the 1.5x interquartile range. Letters denote significant Tukey contrasts, with α < 0.05.

Iron was by far the most common metal across all samples and liming did not restrict its availability (i.e. dissolved concentration in pore water phase: paired t-test: *t* = 0.15, *df* = 29, *P* = 0.88; mean [95% CI] iron availability for C = 77.17[31.92, 122.41] and L = 81.04[50.88, 111.20] mg^-1^ L pore water). As a consequence, total metal availability did not differ between treatments (treatment effect: *χ^2^_1_* = 0.9645, *P* = 0.33; Fig. 2b). However, the availability of non-ferrous metals was significantly lower in soils that had received a single dose of hydrated lime (treatment effect: *χ^2^_1_* = 3.96, *P* = 0.047; Fig. 2c). As a likely result of differing metal solubilities and co-precipitation (Adriano 2001), the effect of liming varied greatly across metals (Fig. 2a and Figure 2 – source data 1). Notably, while none of the quantified metals was more readily available in lime-treated soils, liming did reduce the level of soluble copper (Cu), zinc (Zn), aluminium (Al) and magnesium (Mg), all of which can be toxic at high concentrations (Gadd and Griffiths 1977).

**Figure 2.**
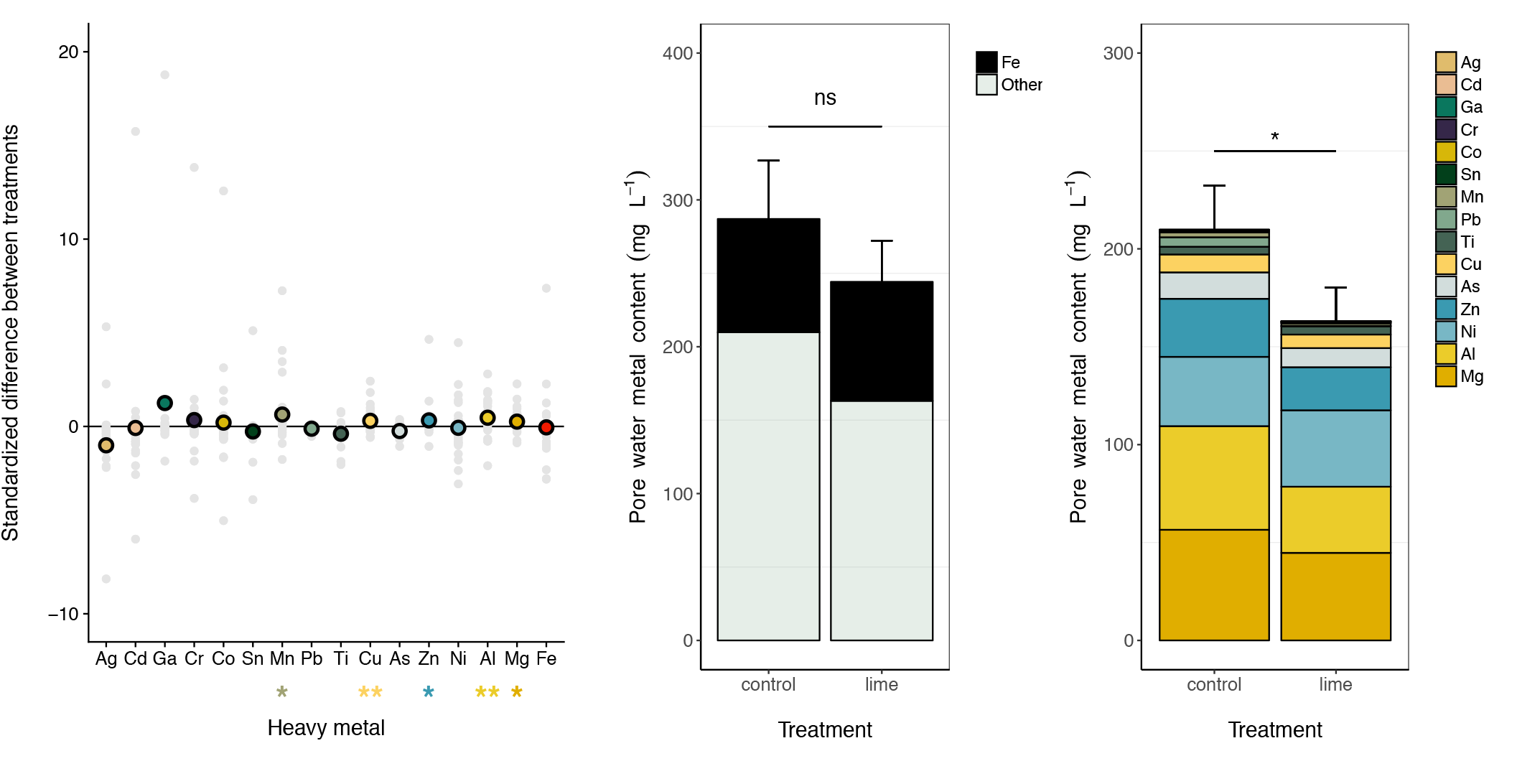
Liming reduces the availability of a subset of non-iron metals**. a.** Plot depicting mean (coloured symbols) and individual (grey symbols) standardized differences between paired soil microcosms, where positive values indicate a reduction in metal availability (i.e. dissolved metal concentration in pore water phase) in limed versus control soils. Stars summarize the significance of this lime effect, estimated using one-tailed t-tests corrected or multiple testing (^**^ *P* < 0.05 and ^*^ *P* < 0.1). Note that a subset of metals (As, Ga, Pb, Sn and Ti) was not included in the statistical analyses as the majority of samples contained undetectable levels. Bar plots demonstrating **(b)** a non-significant effect of liming on total metal availability (mean ± 1 S.E.), partitioned into iron (black) and non-iron (mint green) metals and **(c)** a significant reduction in total non-iron metal availability (mean ± 1 S.E.) n limed soils, partitioned into individual metals, which are arranged based on their mean across-treatment availability, with magnesium (Mg) being most common. Star denotes significant Tukey contrast, with α < 0.05. A source file containing a table summarizing treatment-specific metal availability is available in Figure 2—source data 1.

### Liming alters microbial community composition and selects against detoxifying siderophores

The majority of the variation in community composition between samples (i.e. weighted Unifrac distance) was accounted for by the natural contamination gradient (PERMANOVA, *F_25_,_50_* = 14.45, partial *R*^2^ = 0.85, *P* < 0.001, Fig. 3). While these large between-community differences masked the effect of liming to a certain extent, the relative abundance of several common phyla – including members of the Acidobacteria, Chloroflexi and Gemmatimonadetes – did change in response to lime addition (Figure 3 – source data 1). In agreement, our analysis confirms that different taxa were favoured across different treatments (PERMANOVA, *F_2, 50_* = 6.20, partial *R*^2^ = 0.029, *P* < 0.001): when decomposed into multiple pairwise comparisons, community composition differed significantly between liming and the control (Bonferroni-corrected *P* = 0.016), and liming and the ancestral community (Bonferroni-corrected *P* < 0.001), but not between the control and ancestral community (Bonferroni-corrected *P* = 0.408).

**Figure 3.**
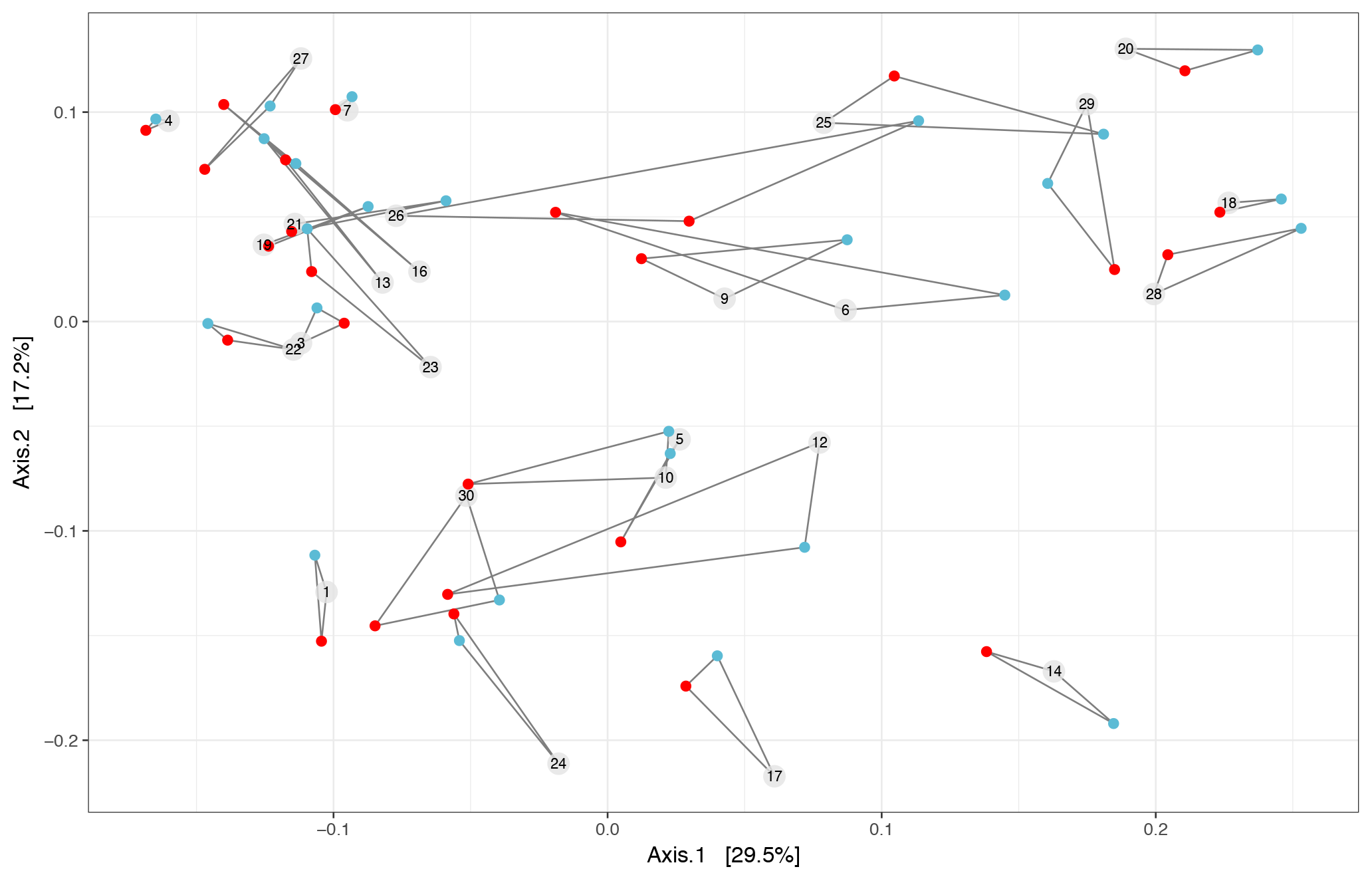
Principal Coordinate Analysis (PCoA) plot based on weighted Unifrac distance of microbial ASVs between communities. The percentage of variation explained is shown on each axis, calculated from the relevant eigenvalues. Communities with the same community origin are joined with straight lines. Although community origin drives most of the variation in community composition, the addition of lime (red circles) also significantly changes community composition compared to the ancestral (labelled in numbered grey circle) and control (blue circles) communities. Communities 2, 8, 11 and 15 were not included in our analyses as DNA yield was not of sufficiently high quality for amplicon sequencing. A source file depicting the effect of liming on the relative abundance of common phyla is available in Figure 3 – source data 1.

Siderophore production by individual clones (*n =* 2016) was assayed by quantifying extracellular iron-chelation activity *in vitro* using liquid CAS assays (Schwyn and Neilands 1987). Microbial communities naturally varied in their siderophore production (lmer: random intercept variance of microbial community = 0.15; Fig. 4a), with all ancestral communities containing multiple siderophore-producing clones (Fig. 4c). Crucially, liming strongly selected against siderophore production (treatment effect: *χ^2^_2_* = 2247.6, *P* < 0.001; Fig. 4a-b). Mean per-capita siderophore production was significantly reduced in lime-treated (mean = −0.25, 95% CI = −0.32 – −0.18) compared to ancestral (mean = 0.31, 95% CI = 0.24 – 0.39) or control soils (mean = 0.56, 95% CI = 0.48 – 0.64) (Tukey contrasts for C–T_0_: *z* = 18.85, L–T_0_: *z* = −42.84 and L–C: *z* = −61.60, all *P* <0.001). This pattern was driven by lime selectively favouring non-siderophore-producing clones (Fig. 4c). Control communities showed an increase in siderophore production (Fig. 4), which was likely caused by selection pressures resulting from laboratory conditions.

**Figure 4.**
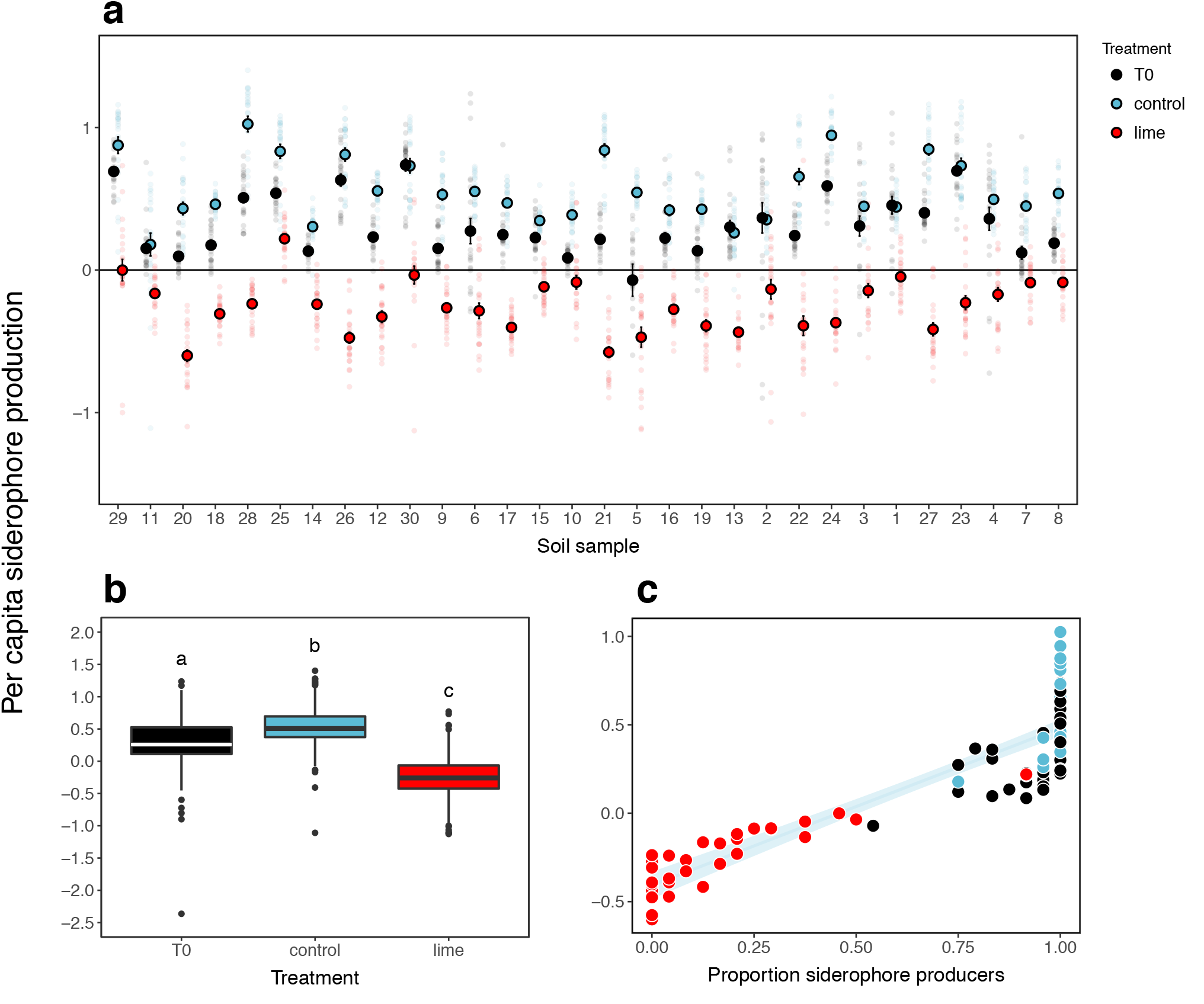
Liming selects against microbial siderophore production. **(a)** Variation in siderophore production across thirty microbial communities as a function of selection regime, where small dots depict raw data (*n* = 24 clones per community) and large coloured dots provide a measure of mean per capita siderophore production ± 1 S.E. in ancestral (black), limed (red) and control (blue) soils, ranked on their initial acidity. **(b)** Boxplot summarizing treatment-specific variation in siderophore production, where different letters denote significant Tukey contrasts (α < 0.05) and **(c)** Plot demonstrating a positive linear relationship between mean per capita siderophore production and the proportion of siderophore producers within each community (*n* = 90). Line and shaded area depict fitted relationship [± 1 S.E.] pooled across treatments: y = - 0.41 [0.04] + 0.89 [0.05] x; *F_1, 88_* 7.8, *P* < 0.001; adjusted *R^2^* = 0.79. Coloured symbols denote the different selection regimes, where black, red and blue are ancestral, limed and control soils, respectively.

## Discussion

Our findings demonstrate that the effect of liming on siderophore production is consistent across a range of metalliferous soil samples despite the fact that: i) microbes can produce multiple siderophores with very different metal affinities (Hider and Kong 2010; Schalk et al. 2011) and ii) ancestral communities varied widely in their initial composition and pH. As liming did not restrict the availability of iron, community-wide adaptive responses to liming were most likely driven by changes in metal toxicity rather than iron scavenging. Intervention practices that buffer the effects of metal toxicity are therefore likely to select against other microbial resistance traits that remediate the environment (e.g. heavy metal sequestration), irrespective of whether these traits primarily benefit the actor or confer cooperative resistance to other members of the microbial community (Niehus et al. 2017).

Our previous work has shown that changes in community-wide siderophore production are primarily shaped by ecological species sorting (Hesse, O’Brien, et al. 2018), although rapid evolutionary change – both via mutation and horizontal gene transfer (Nogueira et al. 2009) – cannot be ruled out. While liming did have a significant effect on community composition, the impact of liming was relatively small compared to that resulting from historical environmental conditions, and liming often selected for different taxa between communities. This would imply that lime addition simply selects against high siderophore-producing microbial taxa, because the marginal costs of siderophore production (Buckling et al. 2007) become greater when not needed for remediation.

Our current and previous findings (Hesse, O’Brien, et al. 2018) demonstrate that siderophore production in soil microbial communities can change rapidly through time, which is perhaps unsurprising as it is largely driven by species sorting. As a consequence, ceasing lime addition will likely result in restoration of microbial siderophore production. This raises the questions as to whether there are any negative consequences associated with chemical remediation on the medium to long-term. While liming reduces soil acidity and metal availability, it does not actually remove toxic heavy metals from the environment. Siderophores or siderophore-producing microbes, on the other hand, can help with heavy metal removal when combined with the use of hyper-accumulating plants (phytoremediation: Mosa et al. 2016; Pilon-Smits 2005; Salt et al. 1995; Sessitsch et al. 2013). Specifically, metal uptake by plants is typically enhanced if metals are bound to microbial siderophores (Dimkpa et al. 2009; Ghosh et al. 2011; Hazotte et al. 2018; Lebeau et al. 2008; Thijs, Langill, et al. 2017; Thijs, Sillen, et al. 2017; Zaidi et al. 2006). While liming of metalliferous soils has been shown to increase plant yield in agricultural settings (Fageria and Baligar 2008), it remains unclear how it might affect plant metal uptake, although trace metal accumulation has been less frequently reported at high (>7) compared to low (<7) soil pH (Sessitsch et al. 2013). While this needs to be addressed experimentally, our findings do suggest that lime-mediated selection against detoxifying siderophores could negate the benefits arising from enhanced biomass accumulation, thereby hampering phytoremediation efficacy.

Liming could also positively impact human- and animal welfare: many pathogenic microbes rely on siderophores to scavenge iron from their hosts (Lam et al. 2018; Niehus et al. 2017). Selection against these so-called ‘virulence factors’ (Holden and Bachman 2015) could potentially alter the extent to which microorganisms behave as opportunistic pathogens in environments commonly treated with lime (e.g. soil, slurry, acid mine drainage, sewage and wastewater). Ironically, other intervention practices specifically aimed at minimizing disease severity may have the opposite effect in the long term. Vaccination can select for faster growing and more virulent parasites when they infect non-immune hosts (Read et al. 2015). Moreover, drugs that block virulence-related social behaviours in *Pseudomonas aeruginosa* selected against naturally occurring avirulent mutants that were otherwise able to invade by exploiting the social behaviours of wild-type bacteria (Köhler et al. 2010). In other words, anti-virulence intervention indirectly selected for more virulent strains.

To conclude, anthropogenic pollution is a major problem worldwide (Bridges and Oldeman 1999; Thijs, Langill, et al. 2017). While there is a pressing need to remediate polluted environments, our findings indicate that liming – a common intervention practice – opposes selection operating on naturally human-beneficial decontamination traits. Microbes facilitate many of the processes mediating ecosystem services, including decomposition and mineralisation, inorganic nutrient cycling, disease causation and suppression, and pollutant removal (Bissett et al. 2013). Understanding the eco-evolutionary consequences of human intervention on microbial traits (e.g. detoxification, virulence, resistance) is key for the engineering of evolutionary resilient microbial communities, having important implications for phytoremediation, with further relevance to global human health and industry (Buckling et al. 2007).

## Methods

### Study site and soil sampling

We collected thirty soil samples along a natural contamination gradient located in a disused polymetallic mine in the Poldice Valley, Cornwall, UK (N: 50°14.10; W: −5°10.23). Soils were collected and processed as described previously (Hesse, O’Brien, et al. 2018), after which soil acidity was quantified on the same day. A small fraction of soil per sample was stored at −80°C for phenotypic assays and DNA extractions; the remainder was used to set up a selection experiment (see below).

### Design of selection experiment

To explicitly test whether liming selects against detoxification in natural microbial communities, we set up experimental microcosms by placing 30 g of soil per sample in duplicate 90 mm Petri dishes. Using a paired design, we imposed two different selection regimes: a single dose of hydrated lime (100 mg of Verve Garden lime dissolved in 5 ml of sterile *dd*H_2_0) was added to half of the paired microcosms and 5 ml of sterile *dd*H_2_O to the remainder. These lime-treated and control microcosms (*n* = 60) were randomly placed and incubated in an environmental chamber at 26°C and 75% relative humidity and were kept moist throughout. After 12 weeks of incubation, we collected samples to (i) quantify soil acidity and heavy metal concentrations, (ii) characterize microbial communities and (iii) prepare freezer stocks for future siderophore assays (see below). Freezer stocks were prepared by vortexing 1 g of soil for 1 min with 6 ml of M9 buffer and sterile glass beads, after which the supernatant was stored at −80°C in a final concentration of 25% glycerol.

### Soil characterization

Soil acidity was quantified prior to and after experimental manipulation by suspending 1 g of soil per sample in 5 ml of 0.01M CaCl_2_, which was then shaken for 30 min and left to stand for 1 h, after which pH was measured using a Jenway 3510 pH meter (Stone, UK) (Hendershot and Lalande 2008). For experimental soil microcosms, we also quantified the concentration of soluble heavy metals (i.e. metals present in interstitial liquid) using the detachment procedure described previously (Govender et al. 2013a; Govender et al. 2013b). Briefly, we suspended 5 g of soil per microcosm in 5 ml of *dd*H_2_O in 50 ml falcon tubes that were gently shaken to disperse soil aggregates and centrifuged for 1 min at 300 rpm to remove solids. 1 ml of supernatant was transferred to Eppendorf tubes and re-spun at 3000 rpm for 3 min to remove final solids. The resulting supernatants were 1:1 diluted in 1% HCl, after which solution chemistry (i.e. concentrations of Ag, Al, As, Cd, Co, Cu, Cr, Fe, Ga, Mg, Mn, Ni, Pb, Sn, Ti and Zn) was determined using ICP-MS. As the overwhelming majority of soil-borne microbes reside within interstitial spaces in pore networks (Coyte et al. 2017; Vos et al. 2013), the presence of soluble metals in interstitial liquid is a good proxy of metal availability and hence toxicity (Bryan et al. 2016).

### DNA extraction and microbial community characterization

To determine how microbial community composition varied across soils we extracted genomic DNA from 250 mg soil per sample (all stored in buffer and C1 solution at −80°C) using the MoBio Powerlyzer PowerSoil© DNA isolation kit (Carlsbad, CA, USA), according to the manufacturer’s protocol with the bead beating parameter set to 4500 rpm for 45 s. Samples were additionally cleaned using the Zymo OneStep PCR Inhibitor Removal Kit following the manufacture’s protocol. The integrity of DNA was confirmed using 1% TAE agarose gels stained with 1x Redsafe DNA Stain (20 000X), yielding a total of 78 high quality DNA samples (*n* = 26 soil samples x 3 treatments; see Fig. 3).

Sequencing of amplicons of the V4 region of the 16S rRNA gene using the Illumina MiSeq 16S Ribosomal RNA Gene Amplicons Workflow was undertaken by the Centre for Genomic Research (Liverpool, UK) using the primers described in Caporaso et al (2011):

F: 5′ACACTCTTTCCCTACACGACGCTCTTCCGATCTNNNNNGTGCCAGCMGCCGCGGTAA3′ R: 5′GTGACTGGAGTTCAGACGTGTGCTCTTCCGATCTGGACTACHVGGGTWTCTAAT3′.

Briefly, 5 μl of DNA entered a first round of PCR with cycle conditions 20 seconds at 95 °C, 15 s at 65 °C, 30 s at 70 °C for 10 cycles, followed by a final 5-minute extension at 72 °C. The primer design incorporates a recognition sequence to allow a secondary nested PCR step. Samples were first purified with Axygen SPRI Beads before entering the second PCR performed to incorporate Illumina sequencing adapter sequences containing indexes (i5 and i7) for sample identification. 15 cycles of PCR were performed using the same conditions as above for a total of 25 cycles. Samples were purified using Axygen SPRI Beads before being quantified using Qubit and assessed using a Fragment Analyzer. Successfully generated amplicon libraries were taken forward. These final libraries were pooled in equimolar amounts using Qubit and Fragment Analyzer data and size selected on the Pippin prep using a size range of 300-600 bp. The quantity and quality of each pool was assessed by Bioanalyzer and subsequently by qPCR using the Illumina Library Quantification Kit from Kapa on a Roche Light Cycler LC480II according to manufacturer’s instructions. The template DNA was denatured according to the protocol described in the Illumina cBot User guide and loaded at 8.5 pM concentration. To help balance the complexity of the amplicon library 15% PhiX was spiked in. The sequencing was carried out on one lane of an Illumina MiSeq at 2×250 bp paired-end sequencing with v2 chemistry. The raw Fastq files were trimmed for the presence of Illumina adapter sequences using Cutadapt version 1.2.1 (Martin 2011), using the option -O 3 (i.e. 3′ end of any reads matching the adapter sequence for 3 bp or more were trimmed). Reads were further trimmed using Sickle version 1.200 with a minimum window quality score of 20. Reads shorter than 20 bp after trimming were removed. If only one of a read pair passed this filter, it was included in the R0 file. We then processed and analysed the trimmed sequence data in R (v 3.4.4) using the packages *‘dada2’* and *‘phyloseq’* (Callahan, McMurdie, et al. 2016; Callahan, Sankaran, et al. 2016). Following the standard full stack workflow (Callahan, Sankaran, et al. 2016), we estimated error rates, inferred and merged sequences, constructed a sequence table, removed chimeric sequences and assigned taxonomy. During processing, forward and reverse reads were truncated between 25-250 and 25-230 nucleotide positions, respectively, due to poor quality scores. Assembled Amplicon Sequence Variants (ASVs) were assigned taxonomy using the Ribosomal Database Project (RDP) (Cole et al. 2013). We estimated the phylogenetic tree using *FastTree* which uses approximate-maximum-likelihood to estimate phylogeny from nucleotide alignments (Price et al. 2010). We then further quality controlled processed sequencing data before analyses were undertaken. We filtered out all reads that had not been assigned to the phylum level, any ASVs that were present in less than 5% of all samples and any reads that were assigned as either cyanobacterial or chloroplast origin. Processing and filtering steps resulted in all the 78 samples remaining for downstream analysis, with a maximum number of reads in a sample of 650,204, minimum of 21,594 and mean of 61,609. Amplicon sequencing data have been deposited as ENA Project PRJEB28850 (https://www.ebi.ac.uk/ena/data/view/PRJEB28850).

### Siderophore assays

For each unique microbial soil community – treatment combination (*n* = 90) we quantified siderophore production by plating out serial-diluted freezer stocks on LB agar plates supplemented with Nystatin (20-μg/ml final concentration), which suppresses fungal growth. Plates were incubated at 28°C for 48 h, after which 24 clones per sample were randomly selected and grown independently in 2 ml of iron-limited CAA medium (5 g Casamino acids, 1.18 g K_2_HPO_4_.3H_2_O, 0.25 g MgSO_4_.7H_2_O per litre, supplemented with 20 mM NaHCO_3_ and 100 μg ml^-1^ human apotransferrin) (Griffin et al. 2004). After 48 h of growth at 28°C, we spun down cultures and assayed supernatants to determine the extent of iron chelation using the liquid CAS assay described by Schwyn and Neilands (1987), modified such that one volume of ddH_2_0 was added to the assay solution (Harrison and Buckling 2005). Siderophore production per clone was estimated using: [1 − (A_*i*_/A_*ref*_)] /[OD_*i*_)], where OD_*i*_ = optical density at 600 nanometer (nm) and A_i_ = absorbance at 630 nm of the assay mixture and A_*ref*_ = absorbance at 630 nm of reference mixture (CAA+CAS; A_*ref*_). We measured siderophore production under common garden conditions to avoid confounding effects of environmental variation *in situ*, causing both differential siderophore induction and soil metal-chelating activities.

### Statistical analyses

The effect of liming on soil acidity and heavy metal content (non-ferrous and total soluble metals) was tested using linear mixed effects models (‘lmer’ function from the ‘lme4’ package: Bates et al. 2014), with random intercepts fitted for individual samples (*n* = 30) to account for soil-specific dependencies. We used a similar approach to test for the effect of liming on mean per-capita siderophore production, with the addition of random intercepts being fitted for individual clones (*n* = 24) nested within each soil community. In general, full models were simplified by sequentially eliminating non-significant terms (*P* > 0.05) following a stepwise deletion procedure, after which the significance of the explanatory variables was established using likelihood ratio tests. In case of significant treatment effects, Tukey contrasts were computed using the ‘*glht*’ function from the R package ‘*multcomp*’ (Hothorn et al. 2008), with α < 0.05. The concentration of soluble metals varied widely, ranging from 0 – 1042 mg per litre of interstitial liquid (Fig. 2). To test how liming affected the availability of rare metals in particular we calculated standardized differences between paired samples by dividing sample-specific metal quantities by the overall mean of each metal. We then used one-tailed t-tests to compare standardized treatment differences to zero, corrected for multiple testing (*‘p.adjust’* with method = *‘fdr’*). We removed some metals (As, Ga, Pb, Sn and Ti) from these analyses as the majority of samples contained undetectable levels (Figure 2 – source data 1).

To compare the composition of the microbial communities, we examined the impact of community origin and treatment (i.e. ancestral sample, liming or control) on the weighted Unifrac distance, which weights the branches of the phylogenetic tree based on the abundance of each ASV. Differences in composition between communities were analysed using the R packages ‘*phyloseq*’ and ‘*vegan*’. Permutational ANOVA tests were run using the ‘*adonis*’ function in the R ‘*vegan*’ package using community origin and treatment as main effects and weighted Unifrac distance as a response term with 9999 permutations. We controlled for the nestedness of the data (treatment within community origin) by limiting the shuffling of each permutation to within samples of the same community origin only. We then did pairwise permutational ANOVAs to better understand which treatments were different from each other by running the same approach with only treatment as a main effect and filtering each treatment out in turn. Significance of *P* values was then determined using Bonferroni correction at a value of *P* < 0.05. We used R Version 3.1.3 for all analyses (R Development Core Team; http://www.r-project.org). Raw phenotypic data are presented in Hesse et al (2018) (https://datadryad.org/review?doi=doi:10.5061/dryad.843814d).

## Acknowledgements

We thank Sharon Uren for running our samples on ICP-MS for soil chemistry quantification.

## Source Data files

Raw data on bacterial phenotypes and soil characteristic are presented in Hesse et al (2018) (https://datadryad.org/review?doi=doi:10.5061/dryad.843814d).

Amplicon sequencing data have been deposited as ENA Project PRJEB28850 (https://www.ebi.ac.uk/ena/data/view/PRJEB28850).

Source Data files relating to Figures 2 (Figure 2 – source data 1) and 3 (Figure 3 – source data 1) have been uploaded as part of the manuscript.

